# Cryogenic X-ray crystallographic studies of biomacromolecules at Turkish Light Source “*Turkish DeLight*”

**DOI:** 10.1101/2022.09.03.506456

**Authors:** Necati Atalay, Enver Kamil Akcan, Mehmet Gül, Esra Ayan, Ebru Destan, Fatma Betül Ertem, Nurettin Tokay, Barış Çakilkaya, Zeliş Nergiz, Gözde Karakadioğlu, Abdullah Kepceoğlu, İlkin Yapici, Bilge Tosun, Nilüfer Baldir, Günseli Yildirim, J Austin Johnson, Ömür Güven, Alaleh Shafiei, Nazlı Eylül Arslan, Merve Yilmaz, Cahine Kulakman, Seyide Seda Paydos, Zeynep Sena Çinal, Kardelen Şabanoğlu, Ayşegül Pazarçeviren, Ayşenur Yilmaz, Başak Canbay, Bengisu Aşci, Esra Kartal, Serra Tavli, Mehmet Çaliseki, Günce Göç, Arif Mermer, Gamze Yeşilay, Sevde Altuntaş, Hiroshi Tateishi, Masami Otsuka, Mikako Fujita, Şaban Tekin, Halilibrahim Çiftçi, Serdar Durdaği, Gizem Dinler Doğanay, Ezgi Karaca, Burcu Kaplan Türköz, Burak Veli Kabasakal, Ahmet Kati, Hasan Demirci

## Abstract

X-ray crystallography is a robust and powerful structural biology technique that provides high-resolution atomic structures of biomacromolecules. Scientists use this technique to unravel mechanistic and structural details of biological macromolecules (e.g. proteins, nucleic acids, protein complexes, protein-nucleic acid complexes, or large biological compartments). Since its inception, single-crystal cryo-crystallography has never been performed in Türkiye due to the lack of a single-crystal X-ray diffractometer. The X-ray diffraction facility recently established at the University of Health Sciences, Istanbul, Türkiye will enable Turkish and international researchers to easily perform high-resolution structural analysis of biomacromolecules from single crystals. Here, we describe the technical and practical outlook of a state-of-the-art home-source X-ray, using lysozyme as a model protein. The methods and practice described in this article can be applied to any biological sample for structural studies. Therefore, this article will be a valuable practical guide from sample preparation to data analysis.

## 1. Introduction

The limits and capabilities of X-ray crystallography have been expanded tremendously together with the availability of enabling X-ray technologies (Maveyraud and Mourey, 2020). Advances in hardware and software development have made X-ray crystallography increasingly more attractive for investigating and understanding the structural dynamics of biomacromolecules (Srivastava et al., 2018). The home-source X-ray diffractometer (XRD) enables the determination of the high-resolution structures of biomacromolecules by employing single-crystal X-ray cryo-crystallography (Smyth and Martin, 2000). The extremely intense and focused X-ray beam and high-quality optics offered by new generation home-source XRDs provide a rapid analysis for the investigation of crystalline materials. Initial data collection and structure determination can be performed at cryogenic temperatures without dependence on time-consuming and costly international synchrotron and X-ray Free Electron Laser (XFEL) facility visits.

The main hardware components of a home-source XRD consist of an X-ray generator, a sample holder, and an X-ray detector (Figure 1a; Table 1). X-ray source material depending on XRD can consist of either dual metal targets such as; Mo/Cu, Cu/Cr, Cu/Co, Cu/Ag, and Ag/Mo; or only single metal Mo and Cu. The incoming collimated X-ray beam is scattered through a single crystal which involves arrays of atoms in a periodic crystalline lattice (Cullity and Weymouth, 1957). Diffraction data that originated from monochromatic X-rays by hitting the crystal is collected on a detector and processed through user-friendly software (Powell, 2017).

**Table 1.** Specifications of X-ray Diffractometer.

|  |  |
| --- | --- |
| **Product name** | XtaLAB Synergy-R |
| **Technique** | Single crystal X-ray diffraction |
| **Core attributes** | Microfocus rotating anode X-ray source diffractometer with hybrid pixel array detector and kappa goniometer |
| **Detector type and parameter** | HyPix-Arc 150° (Curved photon counting detector, Hybrid Photon Counting (HPC) X-ray detector, 147 (W) × 93 (H) × 180 (D) (mm) , 1Ø, 100-240 V, 15 A) |
| **X-ray Source** | PhotonJet-R X-ray source with MicroMax™-007 rotating anode, which incorporates a new mirror design and new alignment hardware. Two target types are available (Mo or Cu) |
| **Goniometer** | Fast kappa geometry goniometer, which allows data collection scan speeds of up to 10°/sec |
| **Accessories** | Oxford Cryostream 800, Oxford Cobra, XtaLAB Flow robotic system, XtalCheck-S, High Pressure Kit |
| **Computer** | External PC, MS Windows® OS |
| **Core dimensions** | 1300 (W) × 1875 (H) × 850 (D) (mm) |
| **Weight** | 600 kg (core unit) |
| **Power requirements** | 1Ø, 200-230 V, 20 A |

**Figure 1.**
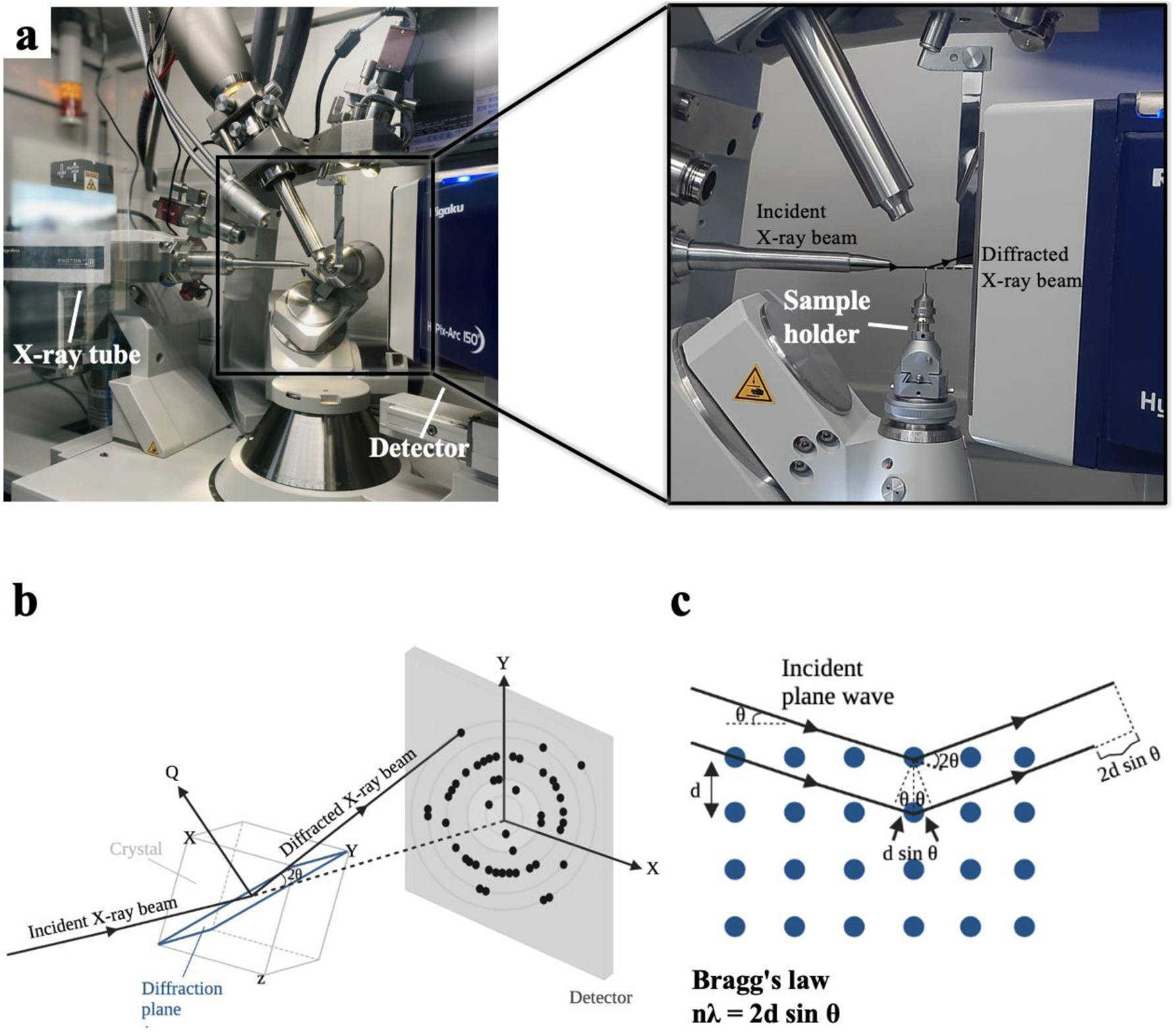
The main hardware components of the Turkish DeLight. **(a)** Schematic of the X-ray diffraction pattern. Incident X-ray beam through the X-ray tube hits the crystal on the sample holder and forms a diffraction pattern based on **(b-c)** Bragg’s law.

Despite recent technological advancements, the phenomenon of X-ray scattering was first defined by Max von Laue, William H. Bragg, and William L. Bragg in the early 20th century (Brügemann and Gerndt, 2004). Based on Bragg’s law, X-ray diffraction is correlated with the characteristic crystal lattice spacing, *d*, with the X-ray wavelength, *λ*, and the angle between the incident and scattered beam, 2*θ* (Figure 1b-1c). When incident X-rays pass through the crystal, based on the principle of Bragg’s law, the X-ray detector converts photon energy to the count rate for recording and processing X-ray signals (Le Pevelen, 2010). Together with the discovery of X-rays in 1895 and the understanding of Bragg’s law, the structural investigation of materials and small molecules had become a milestone for X-ray crystallography (Jaskolski et al., 2014). Single crystal studies have started to highlight the structural and missing properties of macromolecules, which are essential for fundamental biological processes of life (Liebschner et al., 2019; Rathore et al., 2020).

The X-ray diffraction method is very well established and various samples such as small molecules and biomacromolecules can be analyzed (Chapman, 2019; Maveyraud and Mourey, 2020). Nowadays, with expanding computational power, users can determine structures of macromolecules at atomic resolution using high-throughput pipelines (Guven et al., 2021). Macromolecular cryocrystallography experiments were initially carried out at synchrotron X-ray beamlines (Hansen et al., 1990). With the advancement of hard XFELs, ultrafast and bright pulses were achieved and this enabled data collection at ambient temperature while allowing outrunning radiation damage (Chapman et al., 2011). However, limited access to XFEL beamtimes and synchrotrons has been a bottleneck for macromolecular crystallography, limiting the researchers’ speed of molecular characterization. Increasing demand for fully-automated and highly accurate structure determination has paved the way for single-crystal home-source XRDs. Easily accessible XRDs open up new perspectives to the researchers and home source XRDs speed up the whole characterization process. For instance, recently, fast data collection capabilities of home-source XRD enabled rapid ambient-temperature structure determination of SARS-CoV-2 main protease during the COVID-19 pandemic (Kneller et al., 2020).

Here we introduce the Turkish home source XRD, the XtaLAB Synergy Flow system “*Turkish DeLight*” which is equipped with a four-circle kappa goniometer and a universal 6-axis sample handling robot system (UR3) allowing fully automated high-throughput structure determination (Figure 2). High-flux rotating anode X-ray diffractometer and HyPix-Arc 150° detector enable data collection from micron-size crystals with a low detector background noise. The crystal structure determination of the chicken egg lysozyme using a home-source XRD is demonstrated with a step-by-step protocol (Figure 2; Table 2; [CrysAlisPro SOP](https://docs.google.com/document/d/1LEkA-uyhFUmhgO5V-eR4t45Qmv_HVOozKfgUeuS4Eok/edit)). This study is a significant milestone in the Turkish structural biology endeavor as it represents the first high-resolution macromolecule structure determination in which all steps were performed in Türkiye. From crystallization to data collection and processing, all steps were performed without necessitating travel to an international synchrotron light source or an XFEL facility. This has significant implications following a COVID-19 pandemic as fast and efficient drug repurposing studies of SARS-CoV-2 virus were unable to be performed in Türkiye despite the fact that our group successfully obtained two crystal forms of the main protease of this virus (Durdagi et al., 2021). It is an exciting time for Turkish structural biologists that this facility will enable us to perform globally competitive structural biology research and respond to global health and scientific events as they occur.

**Table 2.** Data collection and refinement statistics of lysozyme data collection and structure refinement.

|  |  |
| --- | --- |
| Dataset | Lysozyme |
| PDB ID | 7Y6A |
| **Data collection** |  |
| Beamline | Turkish DeLight Source |
| Space group | P 4<sub>3</sub> 2<sub>1</sub> 2 |
| Cell dimensions |  |
| *a*, *b*, *c* (Å) | 77.35, 77.35, 38.8 |
| $\alpha$, $\beta$, $\gamma$ (°) | 90.00, 90.00, 90.00 |
| Resolution (Å) | 24.46-1.50 (1.78-1.50) |
| *CC*1/2 | 1.000 (0.92) |
| *CC* * | 1.000 (0.97) |
| *R*<sub>merge</sub> | 0.048 (0.44) |
| *R*<sub>meas</sub> | 0.048 (0.46) |
| *I* / σ*I* | 15 (9.8) |
| Completeness (%) | 98.86 (95.99) |
| Redundancy | 21 |
| **Refinement** |  |
| Resolution (Å) | 24.46-1.50 (1.61-1.50) |
| No. reflections | 19182 (1817) |
| *R*<sub>work</sub> / *R*<sub>free</sub> | 0.22/0.25 (0.22/0.25) |
| No. atoms |  |
| Protein | 1006 |
| Water | 134 |
| *B*-factors |  |
| Protein | 17.79 |
| Water | 27.81 |
| Coordinate errors | 0.17 |
| R.m.s deviations |  |
| Bond lengths (Å) | 0.005 |
| Bond angles (°) | 0.720 |
| Ramachandran plot |  |
| Favored (%) | 120 (99.17%) |
| Allowed (%) | 1 (0.83%) |
| Disallowed (%) | 0.00 (0.00%) |

<sup>1</sup>One crystal was used for each dataset. <sup>2</sup>The highest resolution shell is shown in parenthesis.

**Figure 2.**
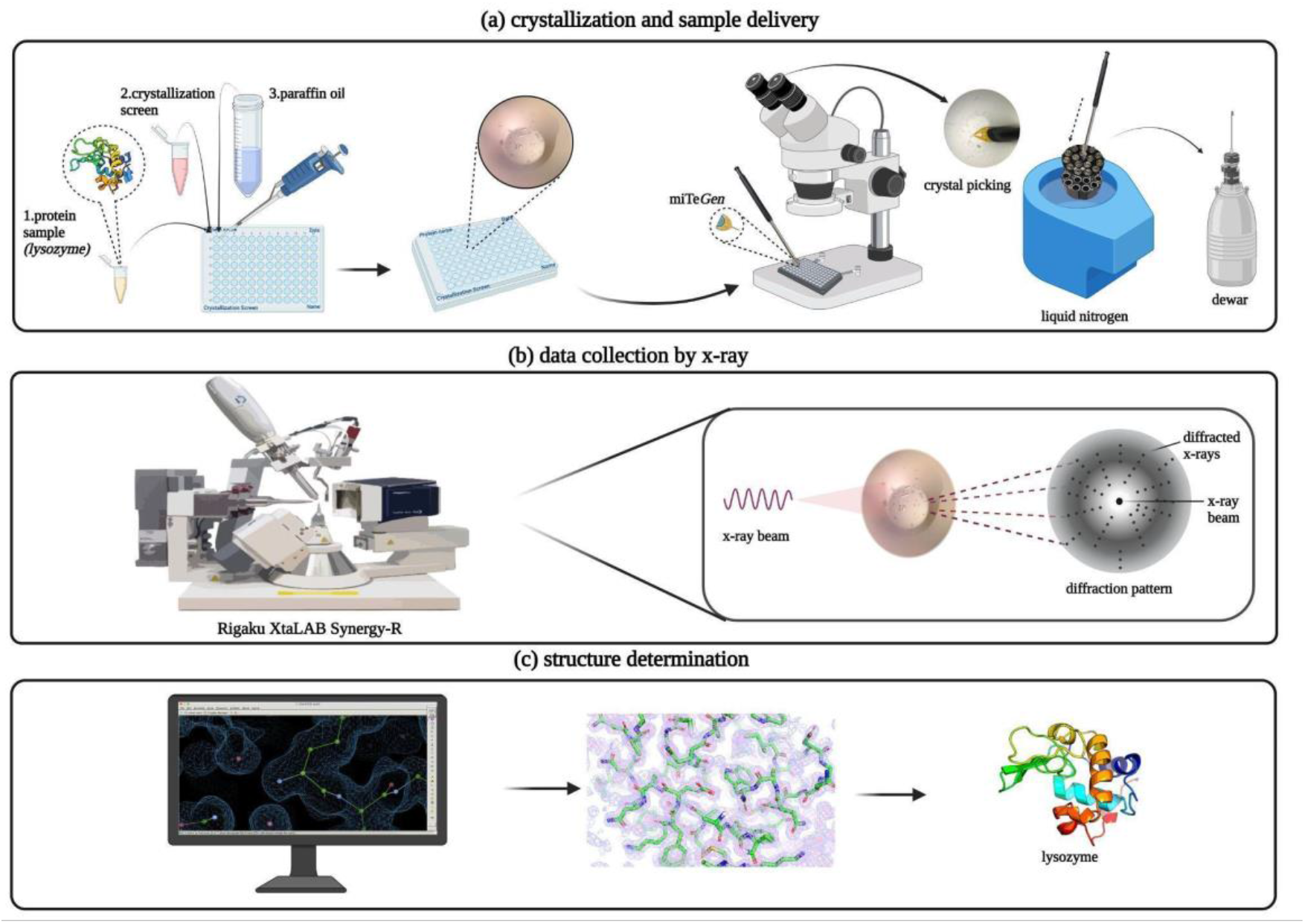
Comprehensive workflow from crystallization to structure determination. **(a)** Crystallization steps of lysozyme and sample delivery; lysozyme crystals are picked by using MiTeGen mounting loops and then the harvested crystals are frozen by plunging into the liquid nitrogen. The puck is transferred to the storage dewar. **(b)** Data collection is performed by using Rigaku’s XtaLAB Synergy Flow X-ray diffractometer. Using the automatic centering feature of the device, the lysozyme crystal at the tip of the MiTeGen loop is centered at the X-ray beam. **(c)** The crystal structure of the lysozyme is determined and refined using *PHENIX* while the model is built in *COOT*. All X-ray crystal structure figures are generated with *PyMOL*.

## 2. Materials and methods

### 2.1 Protein sample preparation and crystallization

Chicken Egg Lysozyme (Calzyme Laboratories, Inc, USA) was prepared from lyophilized powder form by dissolving in distilled water at a final concentration of 30 mg/mL. The protein solution was filtered through 0.22 μm hydrophilic polyethersulfone (PES) membrane filter (Cat#SLGP033NS, Merck Millipore, USA) and stored as 1.0 mL aliquots at −80 °C until crystallization experiments. Sitting drop, micro-batch under oil method was used for lysozyme crystallization (Ertem et al., 2022). 0.83 μL of 30 mg/mL lysozyme solution was mixed with the equal volume of ∼3000 commercially available sparse matrix and grid screen crystallization cocktail solutions in 72-well Terasaki crystallization plates (Cat#654180, Greiner Bio-One, Austria). Each well was sealed with 16.6 μL of paraffin oil (Cat#ZS.100510.5000, ZAG Kimya, Türkiye) and stored at +4 °C until crystal harvesting. Terasaki plates were checked under a compound light microscope for crystal formation and crystals were obtained within 24 hours in most of the crystallization conditions. Overall ∼3000 different crystallization conditions were tested for lysozyme crystallization (Supplementary Table) The best diffracting crystals were grown in 0.09 M HEPES-NaOH pH 7.5, 1.26 M sodium citrate tribasic dihydrate, 10% v/v glycerol (Crystal Screen Cryo (Cat#HR2-122) #38, Hampton Research, USA).

### 2.2 Crystal sample harvesting and delivery

The obtained lysozyme crystals were harvested from the Terasaki crystallization plates by using MiTeGen and microLoops sample pins mounted to a magnetic wand (Garman and Owen, 2006) based on the crystal size (Figure 3) under compound light microscope (Figure 4a-4b). The harvested crystals held by the magnetic wand (Figure 4c) at the tip of the mounting pin were immediately flash-frozen by quickly plunging them in liquid nitrogen. Frozen sample pins were carefully placed on the previously cryo-cooled sample puck (Cat#M-CP-111-021, Mitegen, USA) without removing the crystal from liquid nitrogen (Figure 4d). This step was repeated until the 16-pin puck was completely full. The loaded puck was carefully held using the puck wand (Figure 4e) and transferred to the dewar of the XRD for initial crystal screening and data collection (Figure 4f; please also see [Crystal Picking SOP](https://docs.google.com/document/d/1IDzW37JzQRRZVbT8jye9oKTTqbpQb7wZOqvK10tLb64/edit)). After initial data collection, the remaining crystals in the puck were replaced back to a puck shipping cane (Cat#M-CP-111-065, MiTeGen, USA) and transferred to a CX100 dry liquid nitrogen dewar for long term storage (Cat#TW-CX100, Taylor Wharton, USA).

**Figure 3.**
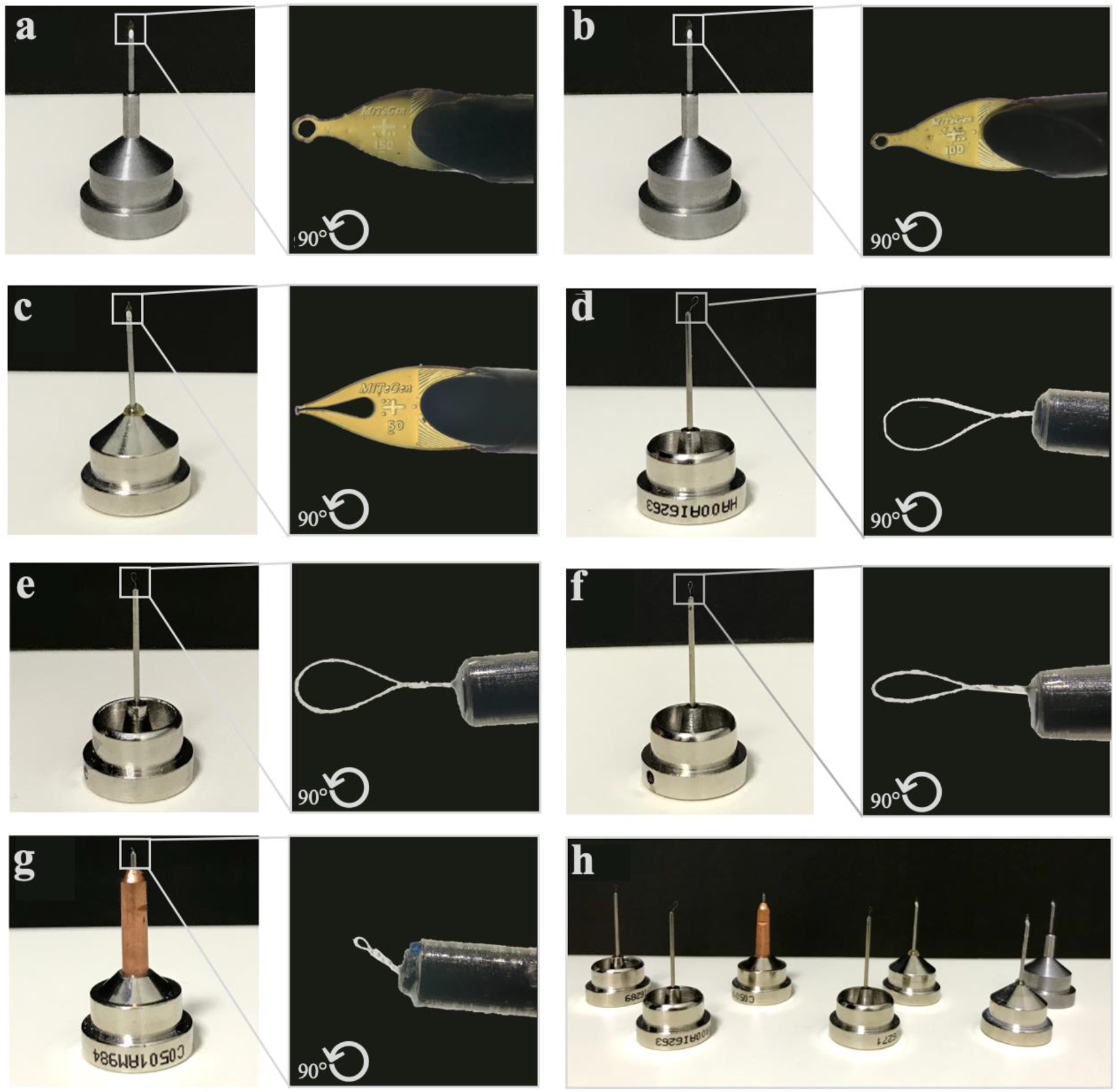
Mounting pins with various mount loops used for crystal delivery. **(a-h)** Different sizes of pins are used based on the crystal size during harvesting.

**Figure 4.**
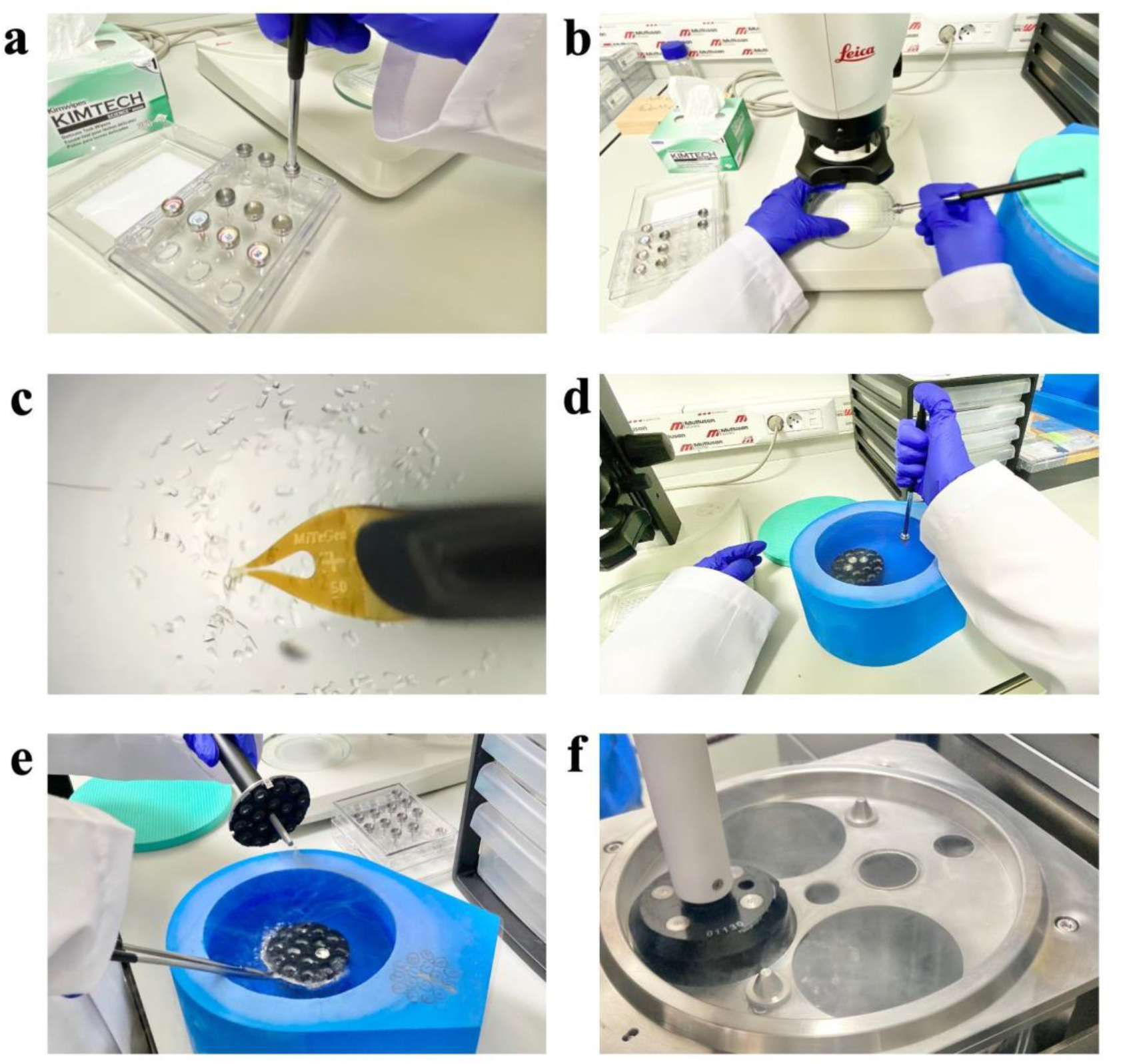
Workflow for sample delivery. **(a)** The loop and cryo-tongs are attached to each other correctly. **(b)** The Terasaki crystallization plate is placed under the compound light microscope and the crystal is located by shifting the plate under the lens. **(c)** The crystal to be collected is picked up with the magnetic tip of the cryopin **(d)** The picked crystal at the tip of the mounting pin is instantly frozen by immersing it in liquid nitrogen, and the pin is inserted into the pre-cooled sample puck in liquid nitrogen. **(e)** Once the crystal freezing is finished, the puck is held by the puck wand, and **(f)** The puck is placed in the dewar of XRD for data collection.

### 2.3 Data collection

Data collection was performed using Rigaku’s XtaLAB Synergy Flow XRD that is equipped with *CrysAlisPro* 1.171.42.35a software (Rigaku Oxford Diffraction, 2021). The sample puck dewar integrated into the device was prefilled and precooled to 100 °K with liquid nitrogen and then the prepared pucks were manually placed in the sample dewar with the puck wand. During the data collection process, to keep the crystals at low temperatures, Oxford Cryosystems’s Cryostream 800 Plus was adjusted to 100 °K and dry air was blown into the sample insertion part of the Intelligent Goniometer Head (IGH). Using the robotic auto sample changer (UR3 as known as Crystal Cracker), the MiTeGen sample pin with a lysozyme crystal was placed on top of the IGH by the help of a robotic arm (Figure 5a). Using the automatic centering feature of the CrysAlisPro, the lysozyme crystal at the tip of the pin was centered at the X-ray beam position. The PhotonJet-R X-ray generator with Cu X-ray source was operated at 40 kV and 30 mA, and the beam intensity was set to 10% to minimize the cross-fire and overlap of the Bragg reflections. Two screening shots were collected with a 2 Å resolution limit set at a 60 mm detector distance, 0.2-degree scan width and 1 second exposure time; a total of 4 frames were taken (Figure 5b). After initial processing, a strategy was created as 1 Å resolution limit, 47 mm detector distance, 0.2-degree scan width, and 1 second exposure time to achieve full data completeness and at least redundancy of 3, with the information obtained from the initial preprocessing of screening images (Figure 5c). A total of 18200 frames were collected within 54 minutes of total data collection time and further used to generate experimental electron density maps extending beyond 1.7 Å.

**Figure 5.**
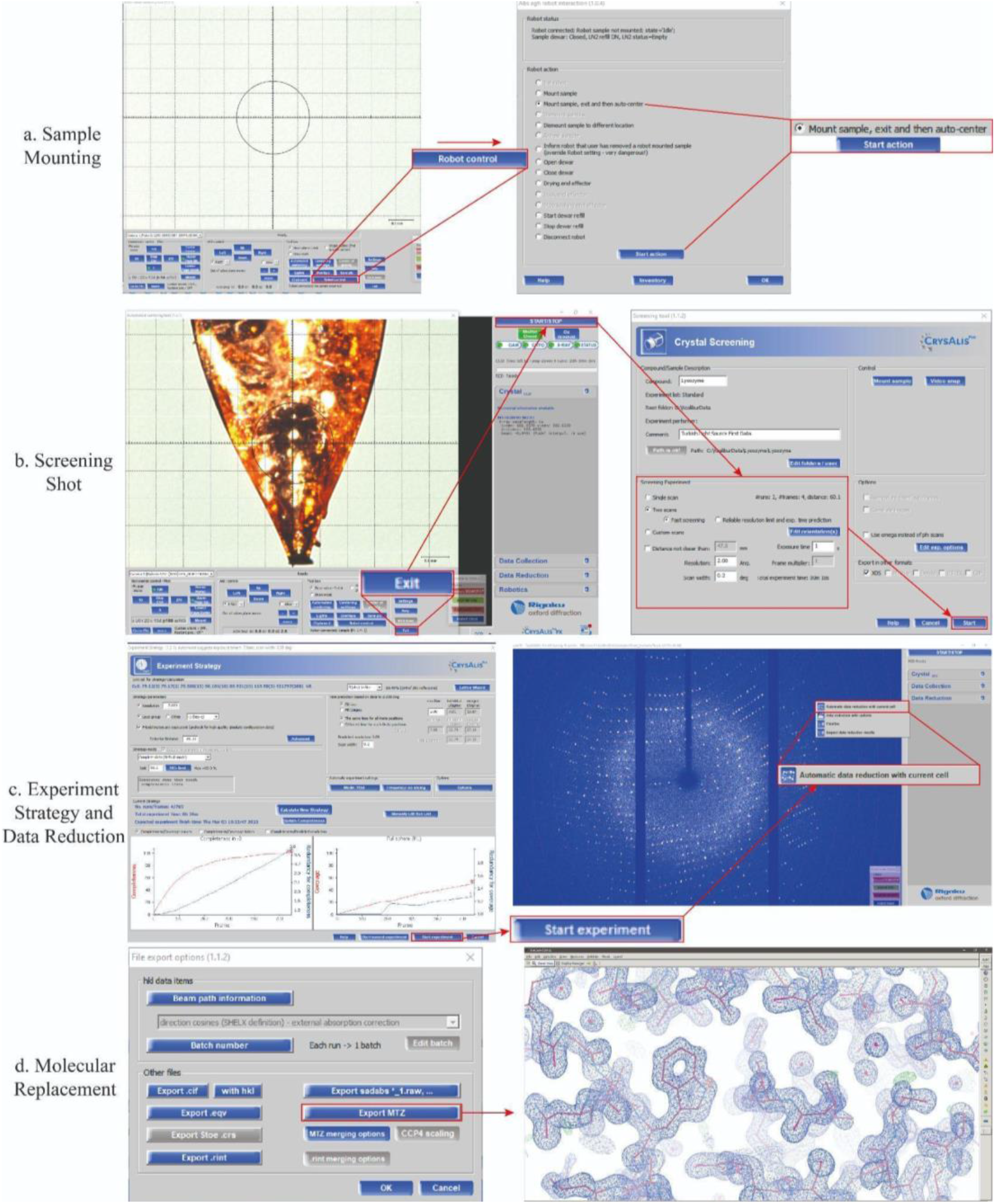
XRD software CrysAlisPro workflow. **(a)** In the control menu of the sample mounting robot, the pin to be placed on the intelligent goniometer head (IGH) is selected. **(b)** Two screening shots with 90° angles provide general information about the crystal by setting the required parameters. **(c)** The necessary parameters are entered, and the data collection strategy is prepared and executed. After the data is collected, an *.hkl file is generated with automated data reduction. **(d)** The *.mtz file is created over the *.hkl file. With the *.mtz file and the template structure PDB: 3IJV, molecular replacement and refinement are performed using *PHENIX*, and the model is built with *COOT*.

### 2.4 Data processing

When the strategy parameters were optimized considering the best diffraction frame (Resolution: 1 Å; Detector distance: 47 mm), and after performing the data collection considering this new strategy, automatic data reduction was started by using the default parameters of the *CrysAlisPro* suite. Moreover, apart from a complete data reduction process in the offline mode, automatic data reduction can be started after batches of 25 frames and updated every 25 frames in the online mode (Figure 5c). The output file is mainly characterized by the *.rrpprof that includes both unmerged and unscaled files. Once data reduction is completed, *.rrpprof file is converted to the *.mtz file using the *.hkl file. In this step, “*CCP4* *.mtz format” can be retrieved through the protein crystallography (PX) window once the *CCP4* package (Winn et al., 2011) is installed. After the data processing is completed, the “Data Reduction” tab in the CrysAlisPro main window provides information on data processing and scaling statistics (Table 2). The data is re-finalized through the “Refinalize/Finalize” button by allowing for the removal of outliers, the re-scaling of frames, and modification of the space group. After determining the export options, the processed data are exported to two *.mtz formats (Figure 5d) that can be used by all PX suites, especially *CCP4* (Winn et al., 2011) and Phenix (Adams et al., 2010).

### 2.5 Structure determination

We determined the crystal structure of lysozyme at cryogenic temperature (100 °K) in space group P4_3_2_1_2 by using the automated molecular replacement program *PHASER* (McCoy et al., 2007) implemented in *PHENIX* (Adams et al., 2010) with the previously published X-ray structure at cryogenic temperature as a search model (Pechkova et al., 2010; PDB ID: 3IJV). Coordinates of the 3IJV were used for the initial rigid-body refinement with the *PHENIX* software package. After simulated-annealing refinement, individual coordinates and Translation/Libration/Screw (*TLS*) parameters were refined. Additionally, we performed composite omit map refinement implemented in *PHENIX* to identify potential positions of altered side chains, and water molecules were checked in *COOT* ([Emsley and Cowtan, 2004](https://www.ncbi.nlm.nih.gov/pmc/articles/PMC8367086/#bib32)), and positions with strong difference density were retained. Water molecules located outside of significant electron density were manually removed. All X-ray crystal structure figures were generated with *PyMOL* and *COOT* (Schrödinger, LLC).

## 3. Results

### 3.1 1.7 Å resolution lysozyme structure determined at Turkish Light Source

We used lysozyme as a model protein to demonstrate a reproducible X-ray data collection from the Turkish DeLight Source and we determined the crystal structure of lysozyme to 1.7 Å resolution at the cryogenic temperature by employing *Turkish DeLight* (Figure 6; Figure 7). The lysozyme structure solved here was aligned to the lysozyme structure (PDB: 3IJV) with an RMSD of 0.44 Å (Figure 8). The Ramachandran statistics for the lysozyme structure (most favored/additionally allowed/disallowed) are 99.17% / 0.83% / 0.00% respectively. The experimentally determined electron density reveals all details within the structure including side chains and water molecules. This well-defined superior electron density map indicates the quality of our data and the small conformational changes were observed as a result of superposition with 3IJV structure.

**Figure 6.**
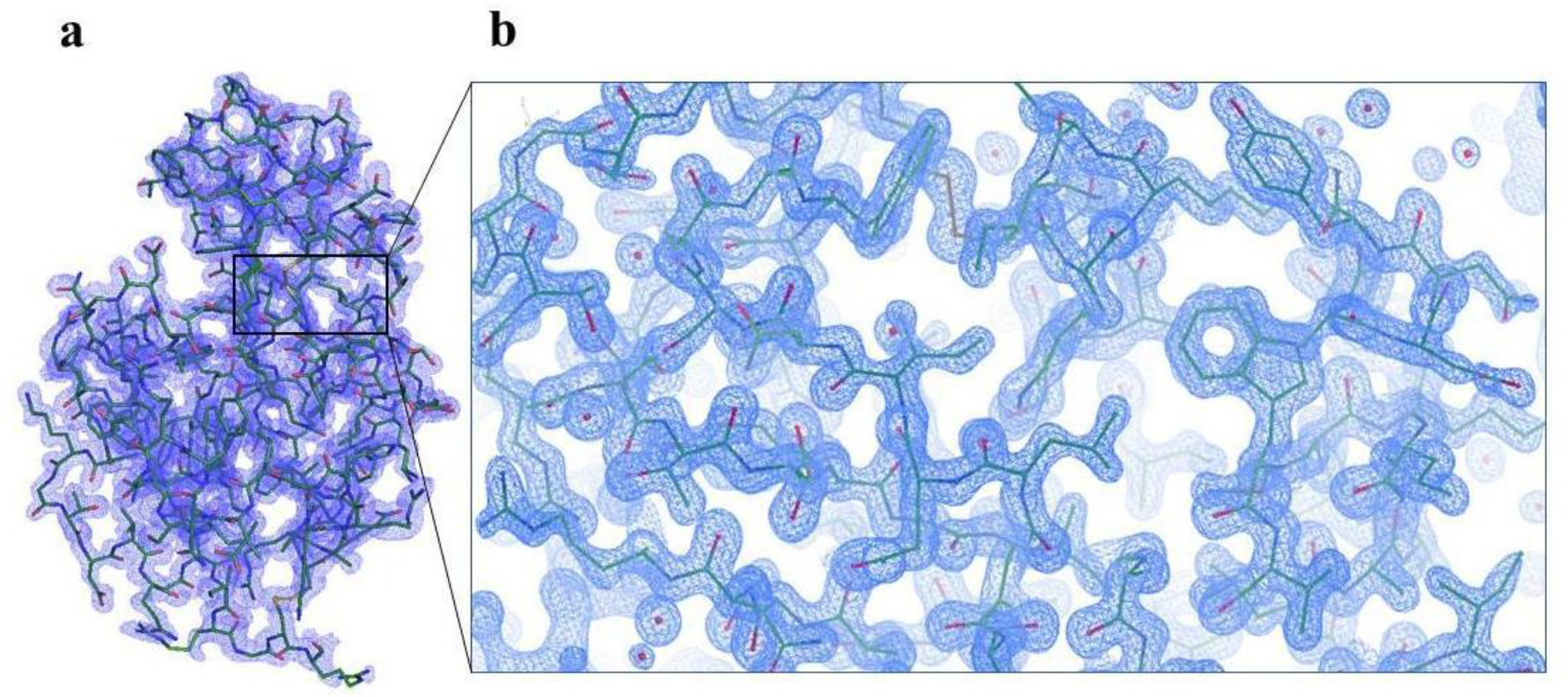
Crystal structure of lysozyme. **(a)** 2Fo-Fc simulated annealing-omit map is colored in blue and shown at 1.0 σ level. **(b)** The crystal structure of lysozyme, is shown in stick representation and colored in slate.

**Figure 7.**
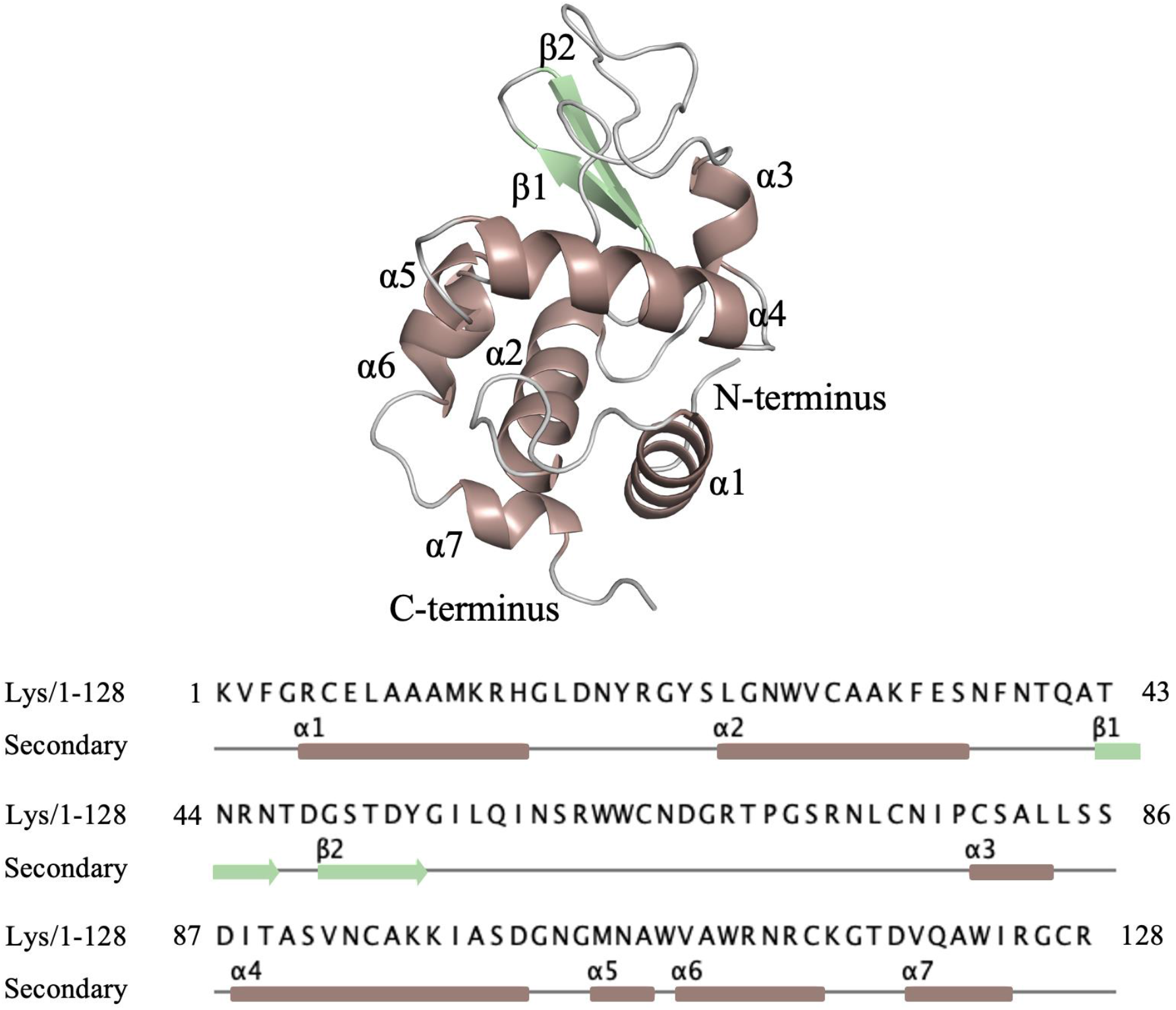
The amino acid sequence of chicken egg lysozyme. The cartoon representation of lysozyme is indicated with secondary structures based on color code, respectively. (Lys: Chicken Egg Lysozyme).

**Figure 8.**
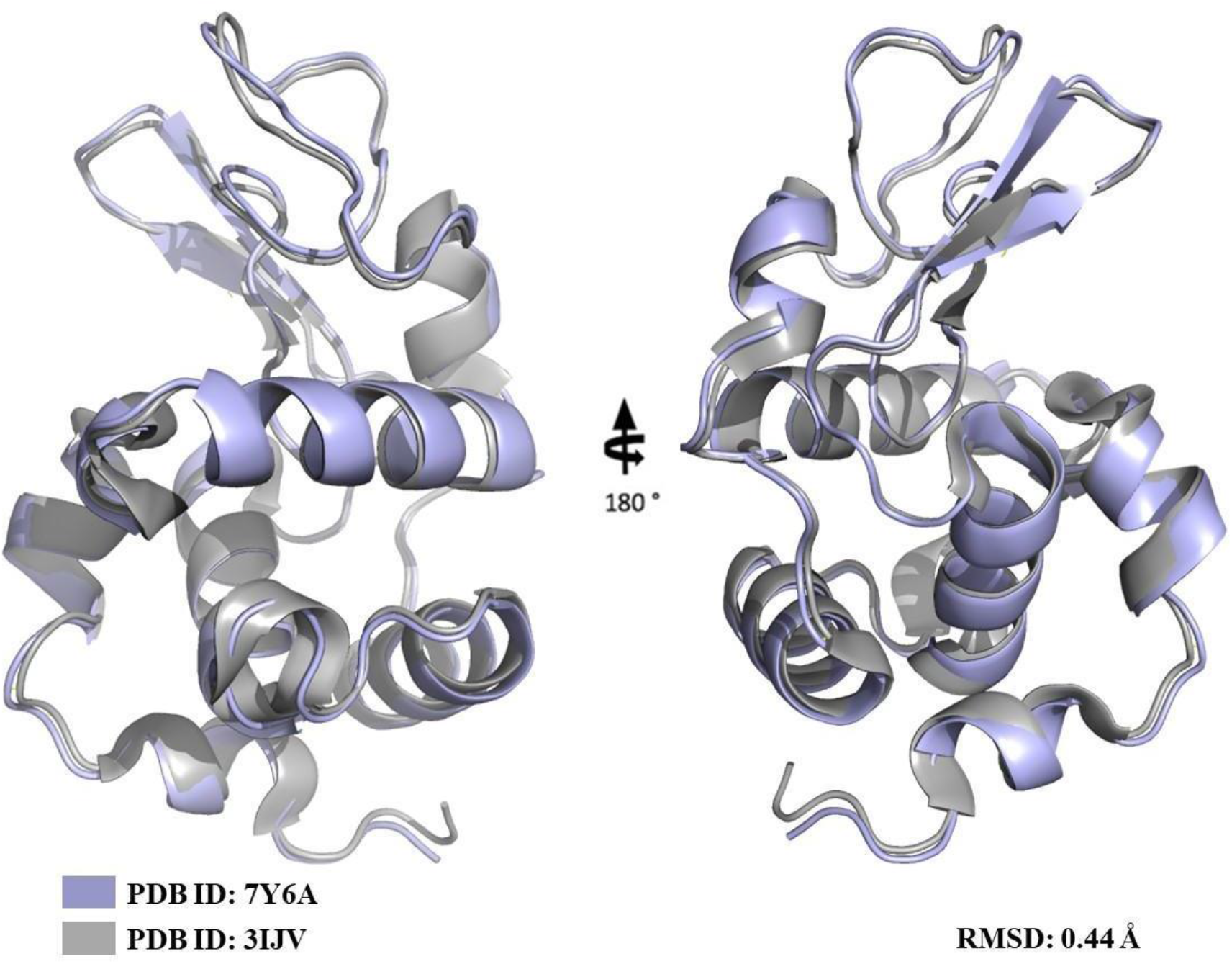
Superposition of the lysozyme structure determined by the data collected at Turkish Light Source and the model lysozyme structure (PDB: 3IJV).

### 3.2 Fully automated data collection and processing

Once picking the best lysozyme crystal based on parameters such as sharp crystal faces, regular shape, and clear rather than opaque, it was flash-frozen in liquid nitrogen. Various crystals were picked and placed until the puck was full and transferred to the dewar of the XRD for the further data collection process (Figure 5; please also see [Crystal Picking SOP](https://docs.google.com/document/d/1IDzW37JzQRRZVbT8jye9oKTTqbpQb7wZOqvK10tLb64/edit)). *CrysAlisPro* offers two different programs: one version is online and supports all data collection and reduction processes; while the other version supports only the data reduction process. Before starting the data collection, the *CrysAlisPro* software should be run with parameters optimized for each experiment before placing the pucks into the dewar. Our study was performed by starting the program with a standard beamstop and IGH, permitting the robot control. We ensured the relative humidity value for the detector was below 10% and that temperature, voltage and current IGH parameters were set to 100 °K, 40 kV and 30 mA through the “CRYO’’ and “XRAY” buttons respectively. Robot control was selected by clicking the “Robotic” button and all processes was performed via the 6-axis UR3 Universal Robot. Once the dewar was filled with liquid nitrogen, all pucks were replaced into the dewar. The inventory zone was adjusted according to the locations of the pucks in the dewar; thus, the crystals were mounted using the auto-centered command of the program via robot action (Figure 5a). Once the auto-centering of a crystal was completed and the relevant parameters were adjusted, screening diffraction was started (Figure 5b). Accordingly, parameters for the beginning of the screen were set as two scans with a 2.0 Å resolution, 0.2-degree scan width, and 0.5 exposure time (Figure 5c). Based on the screening, actual data collection was performed for 54 minutes, establishing the “Go to Strategy” and “Calculate New Strategies” parameters, suggesting the best detector distance is 45 mm and an IUCr limit of 98.5% (Figure 5c). A fully automated space group determination option allows the selection of the best space groups in the Laue Groups. Following the automated data processing, the entire process is finalized by exporting the data as an .mtz file (Figure 5d). Automated molecular replacement was then performed using the *PHASER* implemented in *PHENIX* program package (Figure 5; also see [CrysAlisPro SOP](https://docs.google.com/document/d/1LEkA-uyhFUmhgO5V-eR4t45Qmv_HVOozKfgUeuS4Eok/edit))

### 3.3 XtaLAB Synergy Flow system

*Turkish DeLight* XRD system, configured with XtaLAB Synergy-R, contains high-performance X-ray sources, direct X-ray detection detectors, and 6-axis UR3 Universal Robot that provides the fully- and semi-automated data collection and standardized workflow, providing automated sample mounting/centering, and eliminating potential contamination of the diffractometer. The system is controlled by CrysAlisPro software for further operations, including: sample mounting, queuing, and sorting. Additionally, the HyPix-Arc 150° X-ray detector has Hybrid Photon Counting (HPC) features with a frame rate of 70 Hz. Furthermore, with its 150° curve and high 2θ range, it provides the opportunity to collect data with “Zero Dead Time” without losing any data, and it also guarantees that one can reach high resolutions with its 100-micron pixel size.

### 3.4 CrysAlisPro is a very user friendly X-ray data processing suite

*CrysAlisPro* software provides user-friendly data collection and processing for PX, with an easy-to-use GUI to be operated with manual, fully automated, and semi-automated control. Crystal screening and data reduction can be performed in parallel with fully automated data collection, providing instant experiment feedback using a single integrated package. Unit cells that appear within one or two frames can be investigated over an automated local database or *CellCheckCSD* module during the processing. Similar to the *Ewald3D* tool (3-dimensional diffraction viewer), various tools are available to help the user in identifying and solving problems. In addition, *CrysAlisPro* provides the PX workflows and facilitates remote control of both scientific and technical needs.

## 4. Discussion and conclusion

Many research groups often use their own home source before using synchrotron beamlines to screen and characterize their crystals for getting preliminary data and testing some ligand-binding studies. Therefore, there is a demand for easy to use and efficient XRD infrastructures in which optimum crystal data collection and processing procedures can be performed. These XRD infrastructures should be able to collect high quality and complete dataset by selecting the best crystals for structure determination of macromolecules or ligands. In addition, access and usage of the infrastructure for teaching purposes is also important (Skarzynski, 2013). In this sense, *Turkish DeLight* offers practical advantages to collect rapid and high-quality data in a matter of seconds.

Lysozyme is one of the first enzymes whose structure was determined and best characterized by the X-ray diffraction method. Moreover, lysozyme is widely used in X-ray analysis, unlike other proteins since lysozyme is easy to crystallize and purify from egg white (Sader et al., 2009). We determined the first crystal structure of lysozyme using fully automated data collection and processing via *CrysAlisPro* to create a modern single-crystal XRD pipeline for further analyzing crystals optimizing the PX module over the XtaLAB Synergy-R (Rigaku Oxford Diffraction).

*Turkish DeLight* XRD system is characterized by high-performance X-ray sources, high-throughput HPC X-ray detectors that could be adapted to collect total scattering data and a 6-axis UR3 Universal Robot that provides the fully- and semi-automated data collection. On the other hand, *CrysAlisPro* is a 64-bit compatible user-inspired data collection and processing software with an easy-to-use graphical user interface, which is adaptable for modification and optimization of any PX techniques. Large detectors with high pixel count, more commonly found in synchrotrons, require substantial amounts of memory. Switching to 64-bit with software such as *CrysAlisPro* provides access to more memory and enables easy handling of substantial image sizes and datasets. Additionally, new data image format called *Esperanto* made available at the third-generation synchrotron facility (PETRA III, DESY) is an example of adaptive layouts powered by *CrysAlisPro* for data collecting, processing, and analyzing single-crystal datasets at room temperature (Rothkirch et al., 2013). Likewise, *Esperanto* was also made available for the new diffractometer implemented in a high-resolution inelastic X-ray scattering spectrometer on beamline ID28 at the European Synchrotron Radiation Facility (ESRF) (Girard et al., 2019).

Collectively, this study aims to introduce the first state-of-the-art PX home source XRD in Türkiye. The lysozyme structure determined in this study matches perfectly with the published lysozyme structure (PDB: 3IJV) (RMSD: 0.44 Å), demonstrating that the *Turkish DeLight* offers a high-quality data collection concisely, and presents the user-friendly and easily accessible standard operating procedure (see [Crystal Picking SOP](https://docs.google.com/document/d/1IDzW37JzQRRZVbT8jye9oKTTqbpQb7wZOqvK10tLb64/edit); [CrysAlisPro SOP](https://docs.google.com/document/d/1LEkA-uyhFUmhgO5V-eR4t45Qmv_HVOozKfgUeuS4Eok/edit)) for those who want to collect diffraction data. The future is ultrabright for the next generation of crystallographers in Türkiye.

## Acknowledgement/Disclaimers/Conflict of interest

Authors would like to dedicate this manuscript to the memory of Dr. Albert E. Dahlberg and Dr. Nizar Turker. The authors gratefully acknowledge use of the services and facilities of the Koç University Isbank Infectious Disease Center (KUISCID). H.D. acknowledges support from NSF Science and Technology Center grant NSF-1231306 (Biology with X-ray Lasers, BioXFEL). A.K. acknowledges support from Scientific and Technological Research Council of Türkiye (TÜBİTAK, 2218 - National Postdoctoral Research Fellowship Program under project number 118C476). G.G., M.Ç., and B.V.K. are funded by TÜBİTAK 2232 International Outstanding Researchers Program (Project No: 118C225). This publication has been produced benefiting from the 2232 International Fellowship for Outstanding Researchers Program, 2236 CoCirculation2 program and the 1001 Scientific and Technological Research Projects Funding Program of the TÜBİTAK (Project Nos. 118C270, 121C063 and 120Z520). However, the entire responsibility of the publication belongs to the authors of the publication. The financial support received from TÜBİTAK does not mean that the content of the publication is approved in a scientific sense by TÜBİTAK. Coordinates of the lysozyme structure has been deposited in the Protein Data Bank under accession codes 7Y6A.

