## Supplemental Table for "Cryogenic X-ray crystallographic studies of biomacromolecules at Turkish Light Source “*Turkish DeLight*”"

**Supplementary Table.** Crystallization conditions used for sitting drop, micro-batch (under-oil) crystal screening.

| **Hampton Research** | | |
| --- | --- | --- |
| Natrix (HR2-116) #1-48 | Natrix 2 (HR2-117) #1-48 | Index (HR2-144) #1-96 |
| MembFac (HR2-114) #1-48 | PEG/Ion Screen (HR2-126) #1-48 | PEG/Ion 2 Screen (HR2-098) #1-48 |
| SaltRx 1 (HR2-107) #1-48 | SaltRx 2 (HR2-109) #1-48 | Quik Screen (HR2-221) #1-24 |
| Ionic Liquid Screen (HR2-214) #1-24 | Crystal Screen Cryo (HR2-122) #1-50 | Crystal Screen 2 Cryo (HR2-121) #1-48 |
| Crystal Screen (HR2-110) #1-50 | Crystal Screen 2 (HR2-112) #1-48 | Crystal Screen Lite (HR2-128) #1-50 |
| PEGRx 1 (HR2-082) #1-48 | PEGRx 2 (HR2-084) #1-48 | Grid Screen PEG 6000 (HR2-213) #1-24 |
| Grid Screen Sodium Chloride (HR2-219) #1-24 | Grid Screen Sodium Malonate (HR2-247) #1-24 | Grid Screen Ammonium Sulfate (HR2-211) #1-24 |
| Grid Screen MPD (HR2-215) #1-24 | Grid Screen PEG/LiCl (HR2-217) #1-24 |  |
| **Molecular Dimensions** | | |
| Wizard Classic 1 (MD15-W1-T) #1-48 | Wizard Classic 2 (MD15-W2-T) #1-48 | Wizard Classic 3 (MD15-W3-T) #1-48 |
| Wizard Classic 4 (MD15-W4-T) #1-48 | JCSG-plus™ (MD1-37) #1-96 | HELIX™ (MD1-68) #1-96 |
| MIDASplus™ (MD1-106) #1-96 | NR-LBD™ (MD1-24) #1-48 | NR-LBD™ Extension (MD1-26) #1-48 |
| ProPlex™ (MD1-38) #1-96 | The PGA Screen™ (MD1-50) #1-96 | Morpheus® (MD1-46) #1-96 |
| Structure Screen 1 (MD1-01) #1-50 | Structure Screen 2 (MD1-02) #1-50 | PACT premier™ (MD1-29) #1-96 |
| Stura FootPrint Screen (MD1-20) #1-48 | MultiXtal (MD1-65) #1-48 | MacroSol™ (MD1-22) #1-48 |
| 3D Structure Screen (MD1-13) #1-48 | Wizard Cryo 1 (MD15-C1-T) #1-48 | Wizard Cryo 2 (MD15-C2-T) #1-48 |
| Clear Strategy™ Screen I (MD1-14) #1-24$\times$5 at different pHs 4.5, 5.5, 6.5, 7.5, 8.5 | Clear Strategy™ Screen II (MD1-15) #1-24$\times$5 at different pHs 4.5, 5.5, 6.5, 7.5, 8.5 | Wizard Precipitant Synergy Screen (MD15-PS-T) #1-192 |
| **Jena Bioscience** | | |
| JBScreen Nuc-Pro 1 (CS-181) #1-24 | JBScreen Nuc-Pro 2 (CS-182) #1-24 | JBScreen Nuc-Pro 3 (CS-183) #1-24 |
| JBScreen Nuc-Pro 4 (CS-184) #1-24 |  |  |
| **NeXtal Biotechnologies** | | |
| NeXtal Protein Complex Suite (130715) #1-96 |  |  |
